# An active Unc13A is Reboundless in sleep homeostasis

**DOI:** 10.1101/2025.06.26.661764

**Authors:** Sheng Huang, Marc Escher, Niraja Ramesh, David Toppe, Zhiying Zhao, Yifan Wu, Sha Liu, Alexander M. Walter, Stephan J. Sigrist, Chengji Piao

## Abstract

One of the major characteristics of sleep is homeostatic sleep rebound following sleep loss. While the molecular mechanisms of baseline sleep regulation have been intensively studied, a specific molecular understanding of sleep rebound remains elusive. Here, we show that a constitutively active form of the Munc13-family presynaptic release factor Unc13A, which lacks the inhibitory Ca^2+^/calmodulin interaction domain (Unc13A^WRWR^), dominantly suppressed sleep rebound upon acute sleep deprivation, leading to a nearly complete elimination of recovery sleep (“reboundless”). In contrast, baseline sleep remained largely normal. Through a genetic modifier screen, we found that this dominant “reboundless” phenotype of Unc13A^WRWR^ was rescued by a partial loss of αSnap, a cofactor of NSF required for disassembly and recycling of post-fusion *cis*-SNARE complex. Given that Unc13A promotes fusion-competent *trans*-SNARE complex formation, these findings suggest that sleep rebound may depend on a delicate balance between SNARE complex assembly and recycling. Additionally, we found that expression of a human disease-associated active Unc13A (Unc13A^PL^) variant attenuated baseline and rebound sleep. Since both Unc13A^WRWR^ and Unc13A^PL^ were shown to promote presynaptic release probability (P_r_), we speculate that Unc13A suppresses recovery sleep likely by increasing P_r_ and subsequently enhancing synaptic transmission, probably through elevated *trans*-SNARE formation and efficient *cis*-SNARE recycling. Taken together, our data demonstrate a fundamental role of Unc13A and SNARE dynamics in sleep homeostasis.

## Introduction

Sleep loss resulting from sleep deprivation is prevalent in modern society and is a major driver accelerating a number of human diseases, such as psychological and psychiatric disorders [1-3]. Combating sleep loss and its negative consequences by subsequent compensatory rebound sleep is a common feature of sleep across the animal kingdom and in humans [4, 5]. The “Two-Process” model of sleep regulation proposes that “Process C” controls the circadian timing of sleep and wakefulness, whereas “Process S” regulates homeostatic sleep need [6]. While baseline sleep reflects a fundamental aspect of sleep regulation and function, how a system tracks acute sleep history at the circuit level and subsequently adapts to it remains a central question under intensive investigation in the field [7-9]. Furthermore, the molecular and functional underpinnings for the regulation of sleep rebound without concomitant effects on baseline sleep remains elusive.

Synaptic plasticity is a central feature of sleep function and homeostatic sleep regulation [7, 10-14]. Synaptic structural and functional analyses suggest an increase of synaptic strength coupled with sleep loss, and sleep could reset these changes [7, 11, 13-15]. Indeed, recent studies propose that prolonged wakefulness, such as acute sleep deprivation, leads to the accumulation of sleep need encoded by brain-wide synaptic plasticity via activity changes in specific brain circuits. Furthermore, these synaptic plasticity and circuit activity changes promote a homeostatic sleep rebound in both flies and mammals, thereby establishing a causal link between synaptic plasticity and sleep homeostasis [14, 16]. At the presynapse, where neurotransmitter-filled synaptic vesicles are released, a spectrum of evolutionarily conserved proteins mediate the trafficking, docking, priming and fusion of synaptic vesicles [17-19]. Among these proteins, for fast chemical synaptic transmission, the Munc13-family release factors are essential for vesicle priming by promoting SNARE (soluble N-ethylmaleimide-sensitive factor attachment receptor) complex formation [18-22]. In addition to priming vesicle, Munc13-family proteins have an important post-priming function and “superprime” vesicles to support higher release probability (P_r_) [23]. Mutagenesis and genetic studies suggest an autoinhibitory mechanism of Munc13-family proteins by their regulatory domains such as the polyE motif, Ca^2+^/calmodulin interaction domain (CaM), C1 and C2B domains [24-28]. Furthermore, disrupting these regulatory domains activates Munc13 by relieving the autoinhibition of its MUN domain and enhances synaptic transmission by increasing release probability (P_r_) [24, 25, 27, 29], probably by promoting vesicle superpriming [23] or stabilizing primed vesicles [24]. In *Drosophila*, sleep loss leads to an accumulation of the ELKS-family presynaptic active zone core scaffold protein Bruchpilot (BRP) and the Munc13-family release factor Unc13A [11, 14, 30], which in turn promotes sleep homeostasis [14]. However, how changes in neurotransmitter release properties regulate baseline sleep and sleep homeostasis remains unclear.

In this report, we employed the intensively studied *Drosophila* sleep behavior [31-33], and utilized well-established genetic models of increased P_r_ and consequently enhanced synaptic transmission [24, 27, 34], to dissect the potential role of presynaptic release properties in sleep and sleep homeostasis. We show that activating Munc13-family release factor Unc13A by disrupting its Ca^2+^/calmodulin interaction domain (CaM) (Unc13A^WRWR^) dominantly and nearly completely eliminated sleep rebound upon acute sleep deprivation (“reboundless”). A genetic screen for suppressors of the “reboundless” phenotype highlighttp an important role of α soluble N-ethylmaleimide-sensitive factor attachment protein (αSnap) in sleep homeostasis. Given that Unc13A promotes *trans*-SNARE formation and that αSnap is important for *cis*-SNARE disassembly and recycling [18, 20-22, 35], we propose that Unc13A^WRWR^ promotes P_r_ and suppresses sleep rebound by accelerating the turnover rate of SNARE recycling. Consistently, the human disease-associated Unc13A^PL^ variant, which was shown to promote P_r_ and synaptic transmission [26, 34], also suppressed sleep rebound, suggesting a common scenario of active forms of Unc13A in sleep homeostasis. Our data thus established a molecular and functional explanation for the regulation of sleep rebound, emphasizing the essential role of presynaptic release properties such as P_r_ in baseline sleep and sleep homeostasis.

## Results

The Munc13-family presynaptic release factors are highly conserved across species and promote Ca^2+^-dependent synaptic vesicle exocytosis at the presynaptic active zone [17, 19]. *Unc13A* null mutants have severe defects in fast evoked synaptic transmission and locomotion in *C. elegans*, flies and mammals [36-40]. In contrast, Unc13A with site-directed disruption of the Ca^2+^/calmodulin interaction domain (Unc13A^WRWR^) rescues *Unc13A* null mutants and even increases synaptic transmission when compared to wildtype (*wt*) Unc13A^wt^ control by promoting P_r_ in *Drosophila* [24, 27]. Thus, Unc13A^WRWR^ provides a molecular scenario for testing the behavioral relevance of higher P_r_ and enhanced synaptic transmission, and we started by examining its effects on sleep and sleep homeostasis.

### Unc13A^WRWR^ nearly completely eliminates recovery sleep in *Unc13* mutant background

In *Drosophila*, the *Unc13* locus expresses two major isoforms Unc13A and Unc13B, and the Ca^2+^/calmodulin interaction domain (CaM) is specific to the Unc13A isoform [19, 37]. The double point-mutated Unc13A^WRWR^ mutant was generated based on a genomic Unc13A^wt^ P[acman] construct (Figure 1A) [27]. To avoid endogenous Unc13A expression and potential compensatory effects of endogenous Unc13B when either Unc13A^wt^ or Unc13A^WRWR^ is expressed, we expressed two copies of transgenic Unc13A^wt^ or Unc13A^WRWR^ in an endogenous Unc13-deleted mutant background (*f07072*). We first subjected them to sleep analysis and found that baseline sleep was largely normal in Unc13A^WRWR^ flies compared to Unc13A^wt^, with only a slight reduction in daytime sleep and daytime sleep episode duration, accompanied by an increase in sleep latency at Zeitgeber Time 0 (ZT0). Nighttime sleep, which is normally more consolidated than daytime sleep, did not differ between Unc13A^wt^ and Unc13A^WRWR^ (Figure 1B-1F).

**Figure 1.**
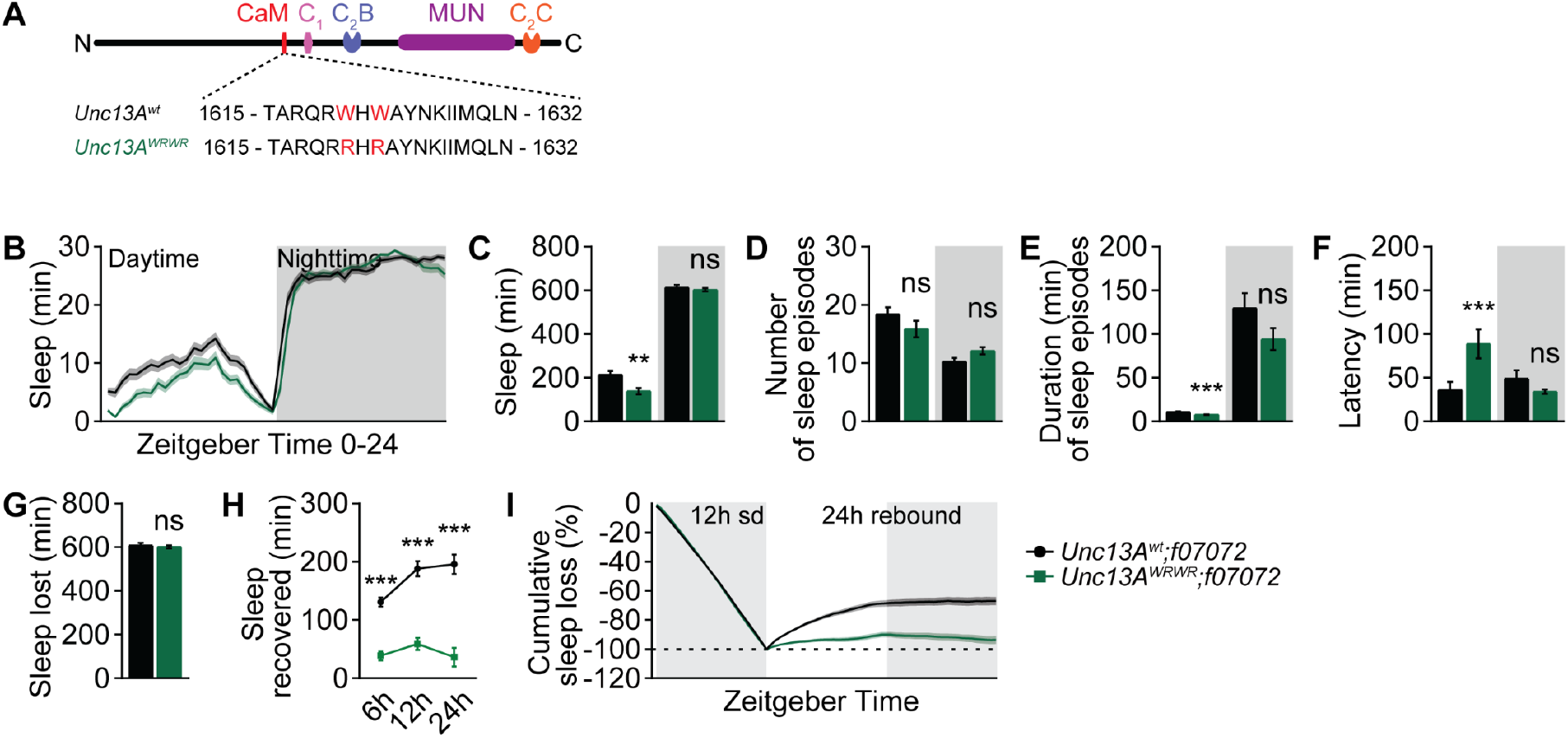
Unc13A^WRWR^ nearly completely eliminates sleep rebound in *Unc13* mutant background. (**A**) Structural domain scheme and the point mutation of the Ca^2+^/calmodulin interaction domain (CaM) of Unc13A (Unc13A^WRWR^). (**B**-**F**) Baseline sleep pattern of *Unc13A*^*wt*^ and *Unc13A*^*WRWR*^ flies in *Unc13*^*f07072*^ mutant background averaged from measurements over 2 days, including sleep profile plotted in 30-min bins (**B**), daytime and nighttime sleep amount (**C**), number and duration of sleep episodes (**D** and **E**), and sleep latency at ZT0 or ZT12 (**F**). n = 61-63. (**G-I**) Sleep deprivation and sleep rebound analysis including sleep loss due to nighttime sleep deprivation (**G**), absolute recovered sleep within 6, 12 and 24 h after sleep deprivation (**H**), and normalized cumulative sleep loss during 12 h nighttime sleep deprivation and subsequent 24 h sleep rebound (**I**). sd, sleep deprivation. n = 61-63. *p < 0.05; **p < 0.01; ***p < 0.001; ns, not significant. Error bars: mean ± SEM.

Besides daily baseline sleep, one major aspect of sleep regulation is the homeostatic sleep rebound following sleep loss, such as acute sleep deprivation [4, 5]. To interfere with baseline sleep for assessing subsequent sleep rebound in these flies, we mechanically deprived the baseline nighttime sleep from ZT12 to ZT24 (Figure 1G), and measured sleep rebound for the next 24 h [14]. In contrast to the largely normal baseline sleep in Unc13A^WRWR^ flies (Figure 1B-F), the sleep rebound of Unc13A^WRWR^ flies following a full night of sleep deprivation was nearly absent when compared to Unc13A^wt^ (Figure 1H and 1I). As Unc13A^WRWR^ is an active form of Unc13A resulting from Unc13A dis-autoinhibition, which triggers an increase of P_r_ [24], these results indicate a unique role of Unc13A activity in specifically regulating sleep rebound without having a major impact on baseline sleep. Furthermore, tuning the efficacy or strength of synaptic transmission by modulating Unc13A activity and P_r_ might be a core mechanism of sleep homeostasis.

Given the unique “reboundless” scenario specific for sleep homeostasis in Unc13A^WRWR^-expressing flies, we would like to alternatively and descriptively refer to this form of Unc13A as “Reboundless” in this study.

### Reboundless Unc13A^WRWR^ dominantly eliminates recovery sleep in *wt* background

Next, we asked if the expression of Unc13A^WRWR^ in the presence of endogenous Unc13A could also effectively suppress sleep rebound by promoting synaptic transmission. At the larval neuromuscular junction, Unc13A^WRWR^ enhances synaptic transmission in *Unc13* mutant background [24, 27]. Here, in *wt* background, we show that *Unc13A*^*WRWR*^ could also promote synaptic transmission (Figure S1), suggesting a dominant effect over endogenous Unc13A. We thus speculated that Unc13A^WRWR^ might dominantly suppress sleep rebound.

We first measured baseline and rebound sleep for flies possessing two genomic copies of either *Unc13A*^*wt*^ or *Unc13A*^*WRWR*^ in *wt* background (Figure 2A-2H). Again, the baseline sleep was largely normal, with a very mild increase in nighttime sleep in *Unc13A*^*WRWR*^ flies compared to *Unc13A*^*wt*^ (Figure 2A-2E), which can be fully deprived by mechanical sleep deprivation (Figure 2F). Importantly, *Unc13A*^*WRWR*^ eliminated sleep rebound in *wt* background (Figure 2F-2H), similar to the effects in *Unc13* mutant background (Figure 1).

**Figure 2.**
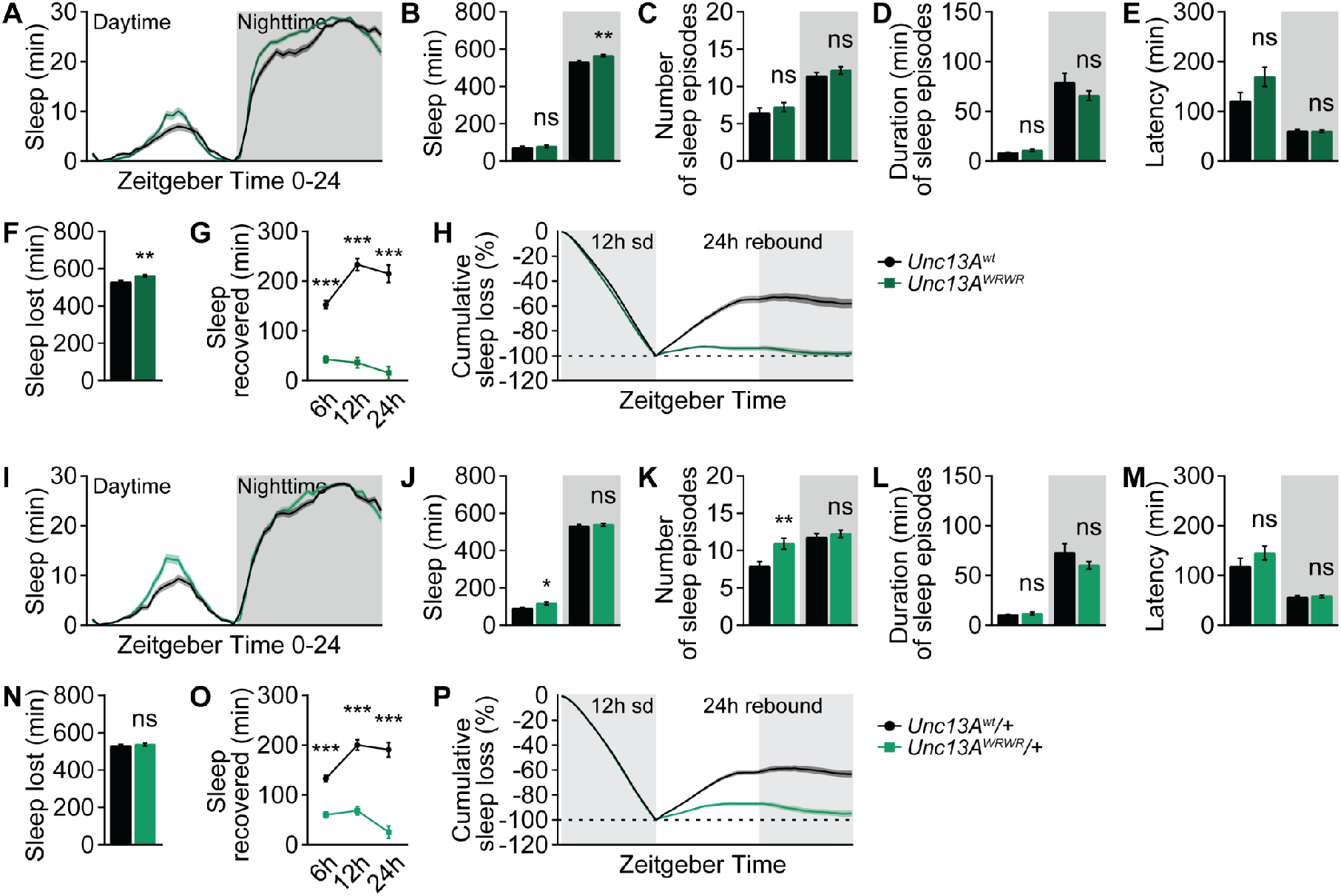
Unc13A^WRWR^ dominantly eliminates sleep rebound in *wt* background. (**A**-**E**) Baseline sleep pattern of *Unc13A*^*wt*^ and *Unc13A*^*WRWR*^ flies in *wt* background averaged from measurements over 2 days, including sleep profile plotted in 30-min bins (**A**), daytime and nighttime sleep amount (**B**), number and duration of sleep episodes (**C** and **D**), and sleep latency at ZT0 or ZT12 (**E**). n = 95-96. (**F-H**) Sleep deprivation and sleep rebound analysis including sleep loss due to nighttime sleep deprivation (**F**), absolute recovered sleep within 6, 12 and 24 h after sleep deprivation (**G**), and normalized cumulative sleep loss during 12 h nighttime sleep deprivation and subsequent 24 h sleep rebound (**H**). sd, sleep deprivation. n = 94-96. (**I**-**M**) Baseline sleep pattern of *Unc13A*^*wt*^/+ and *Unc13A*^*WRWR*^/+ flies in *wt* background averaged from measurements over 2 days, including sleep profile plotted in 30-min bins (**I**), daytime and nighttime sleep amount (**J**), number and duration of sleep episodes (**K** and **L**), and sleep latency at ZT0 or ZT12 (**M**). n = 95-96. (**N-P**) Sleep deprivation and sleep rebound analysis including sleep loss due to nighttime sleep deprivation (**N**), absolute recovered sleep within 6, 12 and 24 h after sleep deprivation (**O**), and normalized cumulative sleep loss during 12 h nighttime sleep deprivation and subsequent 24 h sleep rebound (**P**). sd, sleep deprivation. n = 95-96. *p < 0.05; **p < 0.01; ***p < 0.001; ns, not significant. Error bars: mean ± SEM.

To further test this dominant reboundless phenotype, we introduced a single genomic copy of either *Unc13A*^*wt*^ or *Unc13A*^*WRWR*^ in *wt* background (*Unc13A*^*WRWR*^/+ vs *Unc13A*^*wt*^*/+*, Figure 2I-2P). Similar to the scenario of two genomic copies, the baseline sleep phenotype was consistently very mild, with a slight increase in daytime sleep and daytime sleep episode number (Figure 2I-2M). The suppression of sleep rebound was also very effective (Figure 2N-2P). Thus, these data suggest a specific and dominant role of Unc13A^WRWR^ in synaptic transmission as a potential cause of the reboundless phenotype, highlighting an essential role of fine-tuned synaptic transmission in sleep homeostasis.

### Reboundless Unc13A^WRWR^ suppresses R5 neurons-mediated rebound-like sleep

Recent studies suggest that sleep rebound is regulated by active zone plasticity changes and the activity of ellipsoid body R5 neurons [7, 14]. A brief activation of these neurons for 1 h at ZT0 elicits rebound-like sleep [7]. Given that Unc13A^WRWR^ nearly fully eliminated sleep rebound upon acute sleep deprivation, we wondered if Unc13A^WRWR^ could be able to suppress genetically encoded sleep rebound via activating R5 neurons.

By utilizing an R5-split-Gal4 [7] and the expression of TrpA1 channel [41] to thermogenetically activate R5 neurons at restrictive 32°C for 1 h, we were able to trigger a rebound-like sleep (Figure S2A and S2B), as previously reported [7]. Consistently, baseline sleep at permissive 22°C was again largely normal in *Unc13A*^*WRWR*^/+ compared to *Unc13A*^*wt*^/+, with a very mild increase of both daytime and nighttime sleep, coupled by an increase in daytime sleep episode number and sleep latency at ZT12 (Figure S2C-S2G). We then conditionally activated R5 neuron in *Unc13A*^*WRWR*^/+ compared to *Unc13A*^*wt*^/+ at ZT0 for 1 h, and measured sleep during 1 h R5 neurons activation and rebound-like sleep for the next 23 h. Similar to mechanical sleep deprivation (Figure 2I-2P), *Unc13A*^*WRWR*^/+ flies exhibited significantly reduced rebound-like sleep (Figure S2H and S2I), suggesting a general molecular mechanism of Reboundless Unc13A^WRWR^ in specifically regulating sleep homeostasis.

### Genetic screen for modifiers of the dominant rebound sleep phenotype of *Unc13A*^*WRWR*^/+

The dominant rebound phenotype of *Unc13A*^*WRWR*^/+ flies (Figure 2I-2P) provides an opportunity for genetic modifier screening. As Unc13A^WRWR^ is an active form of Unc13A, of which the MUN domain mediates synaptic vesicle priming and fusion by promoting fusion-competent *trans*-SNARE formation [18, 20, 21], we performed a genetic screen by focusing on key players in synaptic plasticity, presynaptic vesicle trafficking, *trans*-SNARE formation and post-fusion *cis*-SNARE disassembly.

We introduced each heterozygous candidate mutation to either *Unc13A*^*wt*^/+ or *Unc13A*^*WRWR*^/+ background, and measured the difference in baseline and rebound sleep (“Δ baseline” and “Δ rebound”) between *Unc13A*^*wt*^/+ and *Unc13A*^*WRWR*^/+ (Figure 3A-3D). Among all the tested candidates, *fife*/+ [42] did not affect baseline and rebound sleep of *Unc13A*^*WRWR*^/+ flies compared to *Unc13A*^*wt*^/+ (Figure 3E), while *syt1*/+ [43] suppressed rebound in *Unc13A*^*wt*^/+ background and did not affect the rebound phenotype of *Unc13A*^*WRWR*^/+ (Figure 3F). A few synaptic vesicle and SNARE-related heterozygous mutations clearly suppressed “Δ rebound” including *sap47* [44], *complexin* (*cpx*) [45] and αSnap [35] (Figure 3G-3I, highlighted in blue in Figure 3B-3D). Specifically, removing a copy of synaptic vesicle protein Sap47 promoted baseline sleep and alleviated the rebound phenotype (Figure 3B-3D and 3G). In addition, *cpx*/+ also mitigated the rebound defects without affecting baseline sleep in *Unc13A*^*WRWR*^/+ flies (Figure 3B-3D and 3H). Most strikingly, *αSnap*^G8^/+ robustly restored sleep rebound in *Unc13A*^*WRWR*^/+ flies, while baseline sleep did not differ (Figure 3B-3D and 3H). Furthermore, the heterozygous *αSnap*^*M4*^ allele almost fully restore rebound sleep of *Unc13A*^*WRWR*^/+ flies to the level of *Unc13A*^*wt*^/+, while promoting baseline sleep specifically in *Unc13A*^*WRWR*^/+ flies (Figure 3B-3D and 3I-3K).

**Figure 3.**
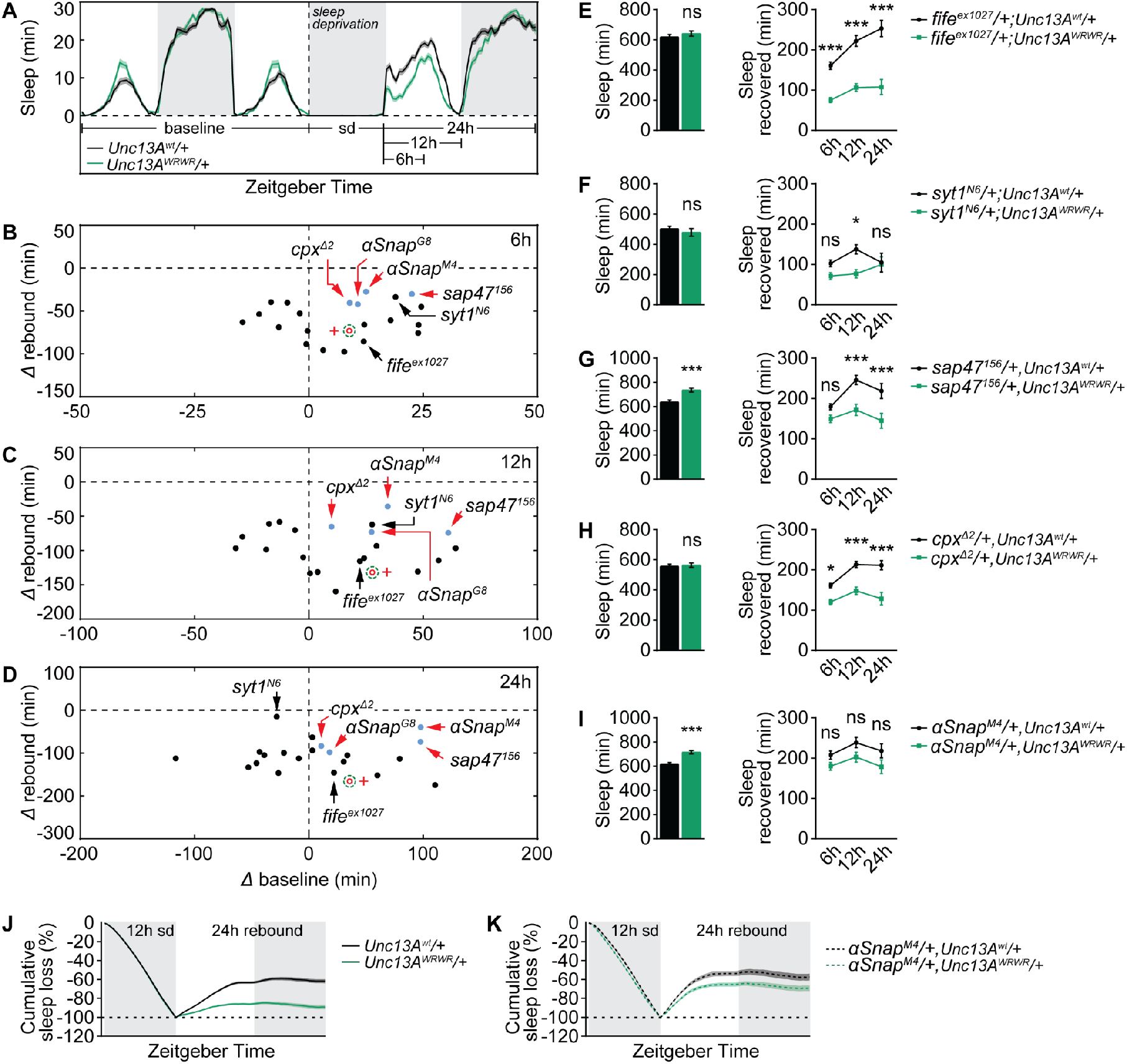
Genetic modifier screen identifies SNARE dynamics in regulating sleep rebound. (**A**) Averaged sleep curve depicting the protocol of mechanical sleep deprivation and sleep rebound. Absolute sleep rebound was analyzed at 3 different time points including 6, 12 and 24 h after sleep deprivation, and the difference between *Unc13A*^*wt*^/+ and *Unc13A*^*WRWR*^/+ is shown as “Δ rebound”. The respective periods of baseline sleep was also analyzed, and the difference between *Unc13A*^*wt*^/+ and *Unc13A*^*WRWR*^/+ is shown as “Δ baseline”. sd, sleep deprivation. (**B**-**D**) Sleep deprivation screen based on the dominant reboundless phenotype of *Unc13A*^*WRWR*^/+. “Δ rebound” is plotted against “Δ baseline” for three different periods of baseline and rebound, including 6 h (ZT0-ZT6, **B**), 12 h (ZT0-ZT12, **C**) and 24 h (ZT0-ZT24, **D**). A few representative heterozygous candidates were labelled and highlighted. Differences between *Unc13A*^*wt*^/+ and *Unc13A*^*WRWR*^/+ in *wt* background are highlighted as “+” and open circles. (**E**-**I**) Baseline 24 h daily sleep and absolute recovered sleep within 6, 12 and 24 h after sleep deprivation for heterozygous candidate mutants including *fife* (**E**, n = 48), *syt1* (**F**, n = 39-48), *sap47* (**G**, n = 72-75), *cpx* (**H**, n = 77-82) and *αSnap* (**I**, n = 63-64). (**J** and **K**) Normalized cumulative sleep loss during 12 h nighttime sleep deprivation and subsequent 24 h sleep rebound for *Unc13A*^*wt*^/+ and *Unc13A*^*WRWR*^/+ either in *wt* background (**J**, n = 173-175) or in *αSnap*^*M4*^/+ background (**K**, n = 63-64). sd, sleep deprivation. *p < 0.05; **p < 0.01; ***p < 0.001; ns, not significant. Error bars: mean ± SEM.

These results suggest that disrupting presynaptic vesicle function and neurotransmitter release can effectively alleviate the rebound phenotype of *Unc13A*^*WRWR*^/+ flies. Given that *αSnap*/+ fully rescued the rebound of *Unc13A*^*WRWR*^/+ flies, we asked if αSnap has any function in regulating sleep and sleep homeostasis. To this end, we tested a hypomorph of *αSnap* (*αSnap*^*f04776*^) and indeed observed a strong increase in both baseline and rebound sleep (Figure S3). Baseline nighttime sleep of *αSnap*^*f04776*^ flies was more consolidated, as indicated by less sleep episodes and longer episode duration (Figure S3A-S3C).

The N-ethylmaleimide-sensitive factor (NSF) and the soluble NSF attachment protein (SNAP) are required for post-fusion *cis*-SNARE disassembly and recycling after vesicle fusion [20, 22]. In *Drosophila*, loss of NSF1 Comatose or SNAP-family member αSnap leads to accumulation of *cis*-SNARE complex [35, 46]. Importantly, αSnap promotes *cis*-SNARE complex disassembly in a dose-dependent manner and *αSnap*/+ animals consistently show elevated levels of *cis*-SNARE complex [35]. Notably, active Unc13A facilitates fusion-competent *trans*-SNARE formation and vesicle priming [18, 20, 21]. These findings support the idea that the rescue of rebound sleep in *Unc13A*^*WRWR*^/+ flies by *αSnap*/+ is mediated through reduced *cis*-SNARE complex disassembly. As a direct consequence, less recycled SNARE components constrain new rounds of *trans*-SNARE formation and vesicle release (also see later detailed Discussion and Figure 4J). This interpretation is further supported by the stronger rebound sleep observed in *αSnap*^*f04776*^ flies (Figure S3).

**Figure 4.**
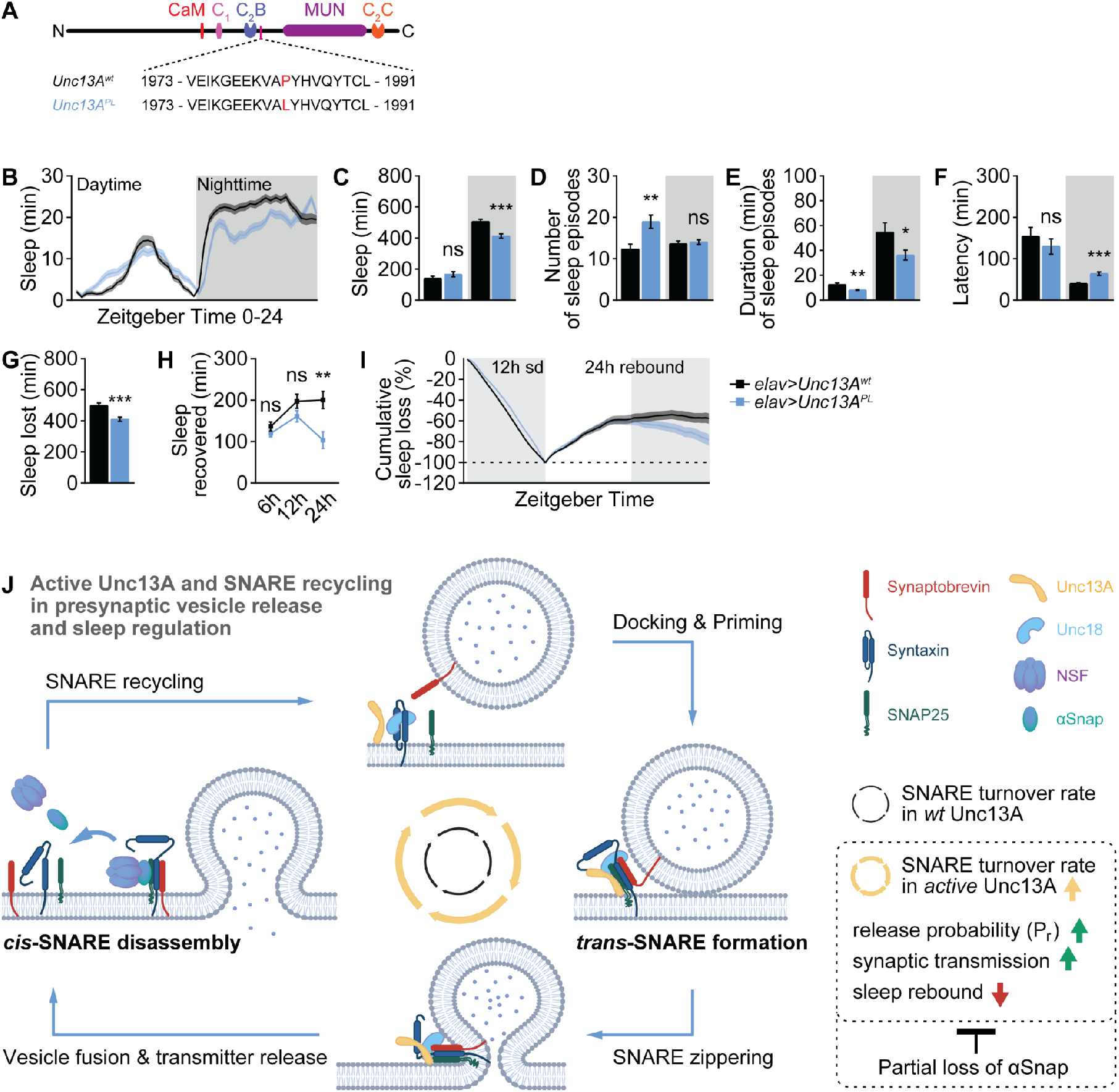
Human disease-associated Unc13A^PL^ variant suppresses baseline and rebound sleep in flies. (**A**) Structural domain scheme of fly Unc13A and its point mutation corresponding to the human disease-associated Unc13A^PL^ variant. (**B**-**F**) Baseline sleep pattern of flies with pan-neuronal expression Unc13A^wt^ or Unc13A^PL^ in *wt* background averaged from measurements over 2 days, including sleep profile plotted in 30-min bins (**B**), daytime and nighttime sleep amount (**C**), number and duration of sleep episodes (**D** and **E**), and sleep latency at ZT0 or ZT12 (**F**). n = 50-55. (**G-I**) Sleep deprivation and sleep rebound analysis including sleep loss due to nighttime sleep deprivation (**G**), absolute recovered sleep within 6, 12 and 24 h after sleep deprivation (**H**), and normalized cumulative sleep loss during 12 h nighttime sleep deprivation and subsequent 24 h sleep rebound (**I**). sd, sleep deprivation. n = 50-55. (**J**) Model for accelerated SNARE turnover rate in promoting P_r_ and synaptic transmission, meanwhile suppressing sleep rebound. Unc13A cooperates with Unc18 to facilitate fusion-competent *trans*-SNARE complex formation and vesicle priming and fusion. Post-fusion *cis*-SNARE complex disassembly requires NSF and its attachment protein αSnap. Disassembled SNARE components (Synaptobrevin, Syntaxin and SNAP25) are then recycled for a new round of vesicle release. Compared to *wt* Unc13A (with intact autoinhibition), constitutive active Unc13A elevates *trans*-SNARE formation and vesicle priming and fusion. Partial loss of αSnap attenuates the function of active Unc13A (potentially in both synaptic transmission and sleep homeostasis) likely by reducing SNARE disassembly and turnover rate, resulting in limited available SNARE components required for new rounds of *trans*-SNARE complex formation. *p < 0.05; **p < 0.01; ***p < 0.001; ns, not significant. Error bars: mean ± SEM.

### Active human disease-associated Unc13A^PL^ variant suppresses baseline and rebound sleep

Finally, apart from Reboundless Unc13A^WRWR^ in flies, an Unc13A^PL^ variant identified in a human patient with developmental delay, autism and movement disorder was shown to have higher P_r_ and increased synaptic transmission in mice [26]. The Unc13A^PL^ variant is a single amino acid exchange from Proline to Leucin at the linker region between the C2B and MUN domains [26]. This linker region is highly conserved and has a crucial role in Unc13A autoinhibition [25]. Similarly, in flies (Figure 4A), the expression of Unc13A^PL^ variant significantly promotes synaptic transmission at larval neuromuscular junction [34].

To examine if the activity of Unc13A regulated by different Unc13A inhibitory regulatory domains has consistent effects on sleep and sleep homeostasis, we pan-neuronally expressed the Unc13A^PL^ variant and measured baseline and rebound sleep (Figure 4B-4I). Compared to Unc13A^wt^, the Unc13A^PL^ variant showed a decrease in nighttime sleep accompanied by less sleep consolidation and longer sleep latency (Figure 4B-4F). Notably, similar to Reboundless Unc13A^WRWR^ (Figure 2), the Unc13A^PL^ variant dominantly suppressed sleep rebound (Figure 4G-4I). Thus, increasing P_r_ via different active forms of Unc13A has consistent effects on sleep homeostasis.

In summary, the presynaptic release properties governed by the Munc13-family factor Unc13A play a pivotal role in sleep homeostasis. Moreover, the findings revealed by behavioral sleep approach combined with a genetic modifier screen suggest that Unc13A promotes SNARE dynamics to support elevated P_r_ and subsequently strengthened synaptic transmission (Figure 4J), offering novel insights into the direct downstream molecular mechanisms, which promote synaptic transmission upon relieving Unc13A autoinhibition.

### Discussion

In this study, we describe a dominant “reboundless” phenotype caused by a CaM-binding site mutation in the essential presynaptic release factor Unc13A (Unc13A^WRWR^). Notably, while rebound sleep is nearly abolished, baseline sleep remains largely unaffected in these flies (Figures 1 and 2). This points to a molecular mechanism that selectively regulates rebound sleep without broadly altering baseline sleep, which is uncommon, as most known regulators of sleep rebound also affect baseline sleep in both flies and mammals [47-52]. Given the evolutionary conservation of Unc13A structure and function[17, 19], this mechanism may offer broader insights into how rebound sleep can be modulated independently of baseline sleep.

To further investigate the mechanisms underlying the reboundless phenotype, we performed a genetic screen and found that heterozygous *αSnap* was sufficient to rescue rebound sleep of Unc13A^WRWR^ flies (Figure 3). αSnap cooperates with NSF1 to disassemble post-fusion *cis*-SNARE complexes [35, 46]. The SNARE complex is composed of plasma membrane-associated Syntaxin and SNAP25, as well as vesicle membrane-localized Synaptobrevin (Figure 4J) [18, 20, 22]. *cis*-SNARE complex disassembly plays an important role in the recycling of SNARE components for another round of *trans*-SNARE complex formation and vesicle fusion (Figure 4J) [22, 53]. Consistently, the accumulation of *cis*-SNARE complexes in *αSnap* mutants likely results in limited SNARE recycling and compromised neurotransmitter release. Along this line, active Unc13A has been shown to promote synaptic transmission through its MUN domain by facilitating *trans*-SNARE complex formation, stabilizing priming of vesicles and reducing energy barrier of synaptic vesicle fusion (Figure 4J) [18, 20, 21, 24, 54], which likely requires efficient recycling of SNARE components after vesicle fusion. It is tempting to speculate that compromising SNARE recycling by reducing *cis*-SNARE disassembly in *αSnap*/+ limits the available SNARE components required for elevated synaptic transmission in Unc13A^WRWR^ flies, thereby alleviates the reboundless phenotype. In addition, *in vitro* studies suggest that αSnap also promotes SNARE zippering during vesicle fusion [55-57]. Thus, alternatively, partial loss of αSnap might reduce synaptic transmission and rescue sleep rebound of Unc13A^WRWR^ flies by less vesicle fusion. In conclusion, our data suggest an important role of SNARE dynamics in sleep homeostasis.

Our data revealed that distinct active forms of Unc13A converge on elevating P_r_ to enhance synaptic transmission, suggesting a critical role of P_r_ in sleep rebound. However, how might Unc13A^WRWR^ and Unc13A^PL^ differ in baseline sleep regulation? The difference might stem from the exact structural and functional status of the activated Unc13A, and subsequently the extent of increases in P_r_ and synaptic transmission, which may regulate distinct aspects of sleep. Considering the essential role of SNARE dynamics in regulating both baseline and rebound sleep (Figure S3), we speculate that sleep behavior is shaped by different Unc13A regulatory domains, which receive distinct upstream inputs, for example Ca^2+^ and diacylglycerol signaling, to regulate the efficacy of *trans*-SNARE formation and vesicle fusion [25, 29, 58, 59].

Taken together, we propose that the active Unc13A^WRWR^ as well as Unc13A^PL^ promote *trans*-SNARE complex formation, which is likely coupled with efficient *cis*-SNARE complex disassembly (Figure 4J), to promote P_r_ and synaptic transmission, and suppress sleep rebound. Future work is warranted in dissecting the exact role of SNARE dynamics in regulating sleep rebound in Unc13A^WRWR^ flies. Our current observation that Reboundless Unc13A^WRWR^ nearly eliminates recovery sleep opens a novel and unique perspective for future work on the molecular mechanisms of sleep homeostasis.

## Materials and methods

Key resources table
| Genotype | Source | Identifier |
| --- | --- | --- |
| Antibodies |  |  |
| Rabbit anti-GFP Alexa 488 | Invitrogen | RRID: AB_221477 |
| Biological Samples |  |  |
| *w<sup>1118</sup>* or *wt* | Bloomington *Drosophila* Stock Center | RRID:BDSC_5905 |
| *Unc13A<sup>wt</sup>* | S. Sigrist lab [37] | N/A |
| *Unc13A<sup>WRWR</sup>* or “Reboundless” | S. Sigrist lab [37] | N/A |
| *Unc13A<sup>f07072</sup>* | Bloomington *Drosophila* Stock Center | FlyBase: FBti0042625 |
| *R5-split-Gal4* | M. N. Wu lab [7] | N/A |
| *UAS-TrpA1* | Bloomington *Drosophila* Stock Center | RRID:BDSC_26263 |
| *UAS-mCD8.GFP* | Bloomington *Drosophila* Stock Center | RRID: BDSC_32188 |
| *fife<sup>ex1027</sup>* | K. M. O'Connor-Giles lab [42] | FlyBase: FBal0280546 |
| *syt1<sup>N6</sup>* | Bloomington *Drosophila* Stock Center | FlyBase: FBal0032850 |
| *sap47<sup>156</sup>* | Bloomington *Drosophila* Stock Center | RRID:BDSC_83712 |
| *cpx<sup>Δ2</sup>* | Bloomington *Drosophila* Stock Center | RRID:BDSC_64252 |
| *αSnap<sup>G8</sup>* | Bloomington *Drosophila* Stock Center | RRID:BDSC_5467 |
| *αSnap<sup>M4</sup>* | Bloomington *Drosophila* Stock Center | RRID:BDSC_5465 |
| *αSnap<sup>f04776</sup>* | Bloomington *Drosophila* Stock Center | RRID:BDSC_85684 |
| *elav-Gal4* | Bloomington *Drosophila* Stock Center | RRID:BDSC_458 |
| *UAS-Unc13A<sup>wt</sup>* | S. Sigrist lab [37] | N/A |
| *UAS-Unc13A<sup>PL</sup>* | S. Sigrist lab [34] | N/A |
| Software and Algorithms |  |  |
| Leica Application Suite X | Leica Microsystems | RRID: SCR_013673 |
| ImageJ (Fiji) | http://fiji.sc | RRID: SCR_002285 |
| GraphPad Prism | GraphPad Software | RRID: SCR_002798 |
| MATLAB R2019a | MathWorks | RRID: SCR_001622 |

### Key resources table

#### *Drosophila* stocks and maintenance

Flies were reared under standard laboratory conditions and raised on semi-defined medium (Bloomington recipe) under a 12/12 h light/dark cycle with 65% humidity at 25°C [60]. 3∼4-day-old female flies were used for all experiments. Except for the genetic sleep rebound screen (Figure 3), all fly strains were backcrossed to *w*^*1118*^ (*iso31*, BDSC#5905) background for at least six generations.

### Sleep measurements and sleep deprivation

Sleep experiments were performed exactly as previously reported [14, 60]. Briefly, sleep of single female flies was measured by Trikinetics *Drosophila* Activity Monitors (DAM2) from Trikinetics Inc. (Waltham, MA) in 12/12 h light/dark cycle with 65% humidity at 25°C. 3∼4-day-old single female flies were loaded into Trikinetics glass tubes (5 mm inner diameter and 65 mm length) with 5% sucrose and 2% agar in one side of the tube. Flies were entrained for at least 24 h and their locomotor activity was measured at 1 min frequency. Data from the first ∼24 h were excluded due to entrainment. A period of immobility without locomotor activity counts lasting for at least 5 min was determined as sleep [31].

Mechanical sleep deprivation was performed as previously described [14]. Briefly, the DAM2 monitors were fixed onto a Vortexer Mounting Plate (Trikinetics) on an Analog Multi-Tube Vortexer controlled by a Trikinetics LC4 light controller and acquisition software for sleep deprivation. A pulse of vortex lasting for 1.2 s was applied to the flies randomly with inter-pulse intervals between 0 s and 40 s to fully sleep deprive flies from ZT12 to ZT24 (nighttime). A minimal mechanical force, which was sufficient to efficiently deprive nighttime sleep, was applied so as to avoid injury to flies during sleep deprivation. Sleep during two days baseline, 12 h nighttime sleep deprivation and 24 h sleep rebound was analyzed using the Sleep and Circadian Analysis MATLAB Program (SCAMP) [61]. Absolute sleep rebound (min) gained within 6, 12 and 14 h following sleep deprivation was calculated. Normalized cumulative sleep loss (%) was calculated as the sleep loss at specific time points during sleep deprivation and rebound divided by the total sleep loss at the end of sleep deprivation, and plotted in 30 min bins. Each sleep and sleep deprivation experiment was repeated at least twice.

### Electrophysiology

Single electrode current clamp recordings at *Drosophila* larval neuromuscular junction were performed on muscle 6 in the abdominal segments 2 and 3 of third instar larvae at room temperature as previously described [34]. Larvae were dissected in modified Ca^2+^-free hemolymph-like saline (HL3; 70 mM NaCl, 5 mM KCl, 10 mM NaHCO_3_, 4 mM MgCl_2_, 5 mM HEPES, 5 mM Trehalose, 115 mM Sucrose, pH = 7.2). Recordings were then carried out in 2 ml of HL3 with 0.4 mM CaCl_2_ added. Preparations were visualized under a BX51WI Olympus microscope with a 40× LUMPlanFL/IR water immersion objective (Olympus Corporation, Shinjuku, Tokyo, Japan). Using sharp electrodes filled with 3M KCl, recordings were made from muscle cells with input resistances of ≥ 4 MΩ and initial membrane potentials between -40 and -80 mV. Two cells were recorded per animal. Single evoked excitatory postsynaptic potentials (eEPSPs) were recorded after stimulating the cut end of the segmental motor neuron bundle with 5 V, 300 μs at 0.2 Hz using an S48 Stimulator (Grass Instruments, Astro-Med, Inc., RI, USA). The recording electrodes (30-50 MΩ) were pulled using a Flaming Brown Model P-97 micropipette puller (Sutter Instrument, CA, USA). The stimulation electrodes were pulled using a DMZ-Universal-Electrode puller (Zeitz-Instruments Vertriebs GmbH, Martinsried, Germany), then tip openings were shaped by fire-polishing with a DMF1000 microforge (World Precision Instruments Inc., FL, USA). Recordings were acquired using an Axoclamp 2 B amplifier with HS-2A x0.1 head stage (Molecular Devices, CA, USA) and sampled at 10 kHz with an Axon Digidata 1322 A digitizer (Molecular Devices, CA, USA). Signals were low pass filtered at 1 kHz using an LPBF-48DG output filter (NPI Electronic, Tamm, Germany).

The eEPSP traces were analyzed using custom-written Python scripts utilizing the pyABF package for Python 3.10 [62], as described below. An average trace was generated from 20 eEPSPs traces per cell for single 0.2 Hz stimulation.

Stimulation artifacts of eEPSPs were removed for clarity. Rise time was calculated from the average trace as the time from 10% to 90% of the total amplitude before the peak. Decay constant τ was calculated by fitting a first order decay function to the region of the average trace of the single 0.2 Hz stimulation recording from 60% to 5% of the total amplitude after the peak.

### Genetic screen for modifiers of the rebound phenotype of Unc13A^WRWR^/+

To carry out the genetic modifier sleep screen for the reboundless phenotype of Unc13A^WRWR^/+, we crossed female virgin *Unc13A*^*wt*^ or *Unc13A*^*WRWR*^ flies to candidate mutants to introduce heterozygosity of candidate mutations to either *Unc13A*^*wt*^/+ or *Unc13A*^*WRWR*^/+ flies. *Unc13A*^*wt*^/+ or *Unc13A*^*WRWR*^/+ flies in each heterozygous candidate mutant background were always simultaneously sleep deprived and compared. For each measurement or biological replicate, differences between *Unc13A*^*wt*^/+ and *Unc13A*^*WRWR*^/+ in baseline and rebound sleep (“Δ baseline” and “Δ rebound”) were plotted two-dimensionally for each candidate. At least two biological replicates were performed for each candidate mutant.

### Thermogenetic activation of R5 neurons

Flies with *R5-split-Gal4*-driven thermo-sensitive TrpA1 expression were raised at permissive 22°C and maintained at this temperature before the sleep experiment. 3∼4-day-old single female flies were then loaded onto DAM2 monitors and sleep was recorded. Similar to mechanical sleep deprivation, two days of baseline was acquired at permissive 22°C before activating R5 neurons by increasing the temperature to 32°C for 1 h at ZT0. Temperature then rapidly returned back to 22°C for measuring rebound-like sleep. Cumulative gain of sleep (min) was plotted in 30 min bins. Normalized sleep rebound gained within 6, 12 and 24 h following R5 activation was calculated by dividing sleep gain by the respective baseline sleep.

### Whole-mount brain immunostaining

Whole-mount brain immunostaining was performed as previously reported [60]. 5-day old female *R5-split-Gal4>mCD8-GFP* flies were kept on ice for 1-2 min and then dissected in ice-cold Ringer’s solution (130 mM NaCl, 5 mM KCl, 2 mM MgCl_2_, 2 mM CaCl_2_, 5 mM HEPES, 36 mM Sucrose, pH = 7.3). Dissected brains were fixed immediately in 4% Paraformaldehyde (w/v) in PBS for 30 min on an orbital shaker at room temperature. Samples were washed in 0.7% Triton-X (v/v) in PBS (0.7 % PBST) for 20 min × 3 times, followed by blocking with 10% normal goat serum (v/v) in 0.7 % PBST for at least 2h at room temperature on shaker. After overnight antibody (rabbit anti-GFP conjugated with Alexa 488, 1:500) incubation at 4ºC in darkness, brains were washed in 0.7% PBST for 30 min × 6 times at room temperature. Afterwards, brains were directly mounted on microscope slides in Vectashield and kept in a dark place at 4 ºC before being scanned.

### Confocal microscopy and image analysis

Whole-mount adult brain samples were imaged on a Leica TCS SP8 confocal microscope from Leica Microsystems, and images were obtained using the Leica Application Suite X. Image stacks were processed and analyzed with Fiji (https://fiji.sc/).

### Statistics

Statistics were performed as previously described [14]. GraphPad Prism 6 was used to perform statistics and figures were generated using both Adobe Illustrator and Prism. Student’s t test was used for comparison between two groups and two-way ANOVA with Sidak’s multiple comparisons test was used for multiple comparisons between two different factors (e.g., genotypes vs time).

## Acknowledgments

We thank Mark N. Wu, Kate M. O’Connor-Giles and Bloomington *Drosophila* Stock Center (BDSC) for fly lines. This work was supported by grants from European Research Council (ERC) advanced grant (SynProtect 101097053) and the Deutsche Forschungsgemeinschaft (CRC1315/A08 Project ID 327654276). C.P. was supported by Deutsche Forschungsgemeinschaft (DFG) under FOR5228/RP8 (Project ID 447288260) and NeuroNex (Project ID 436260754).

S.H. was supported by the Leibniz SAW SyMetAge and Charité NeuroCure. NeuroCure is funded by DFG under Germany’s Excellence Strategy - EXC-2049 - 390688087.

## Author contributions

Conceptualization: S.H., and C.P.; Investigation: S.H., M.E., N.R., D.T., Z.Z., Y.W., S.L., A.M.W., S.J.S., and C.P.; Funding acquisition and supervision: S.H., S.J.S., and C.P.; Writing: S.H., and C.P.

## Data availability statement

The authors confirm that all data underlying the findings are fully available without restriction. All relevant data are within the paper and its Supporting Information files. Further information and requests for resources and reagents should be directed to and will be fulfilled by Stephan J. Sigrist and Chengji Piao.

## Figures and figure legends

**Supplementary Figure S1.**
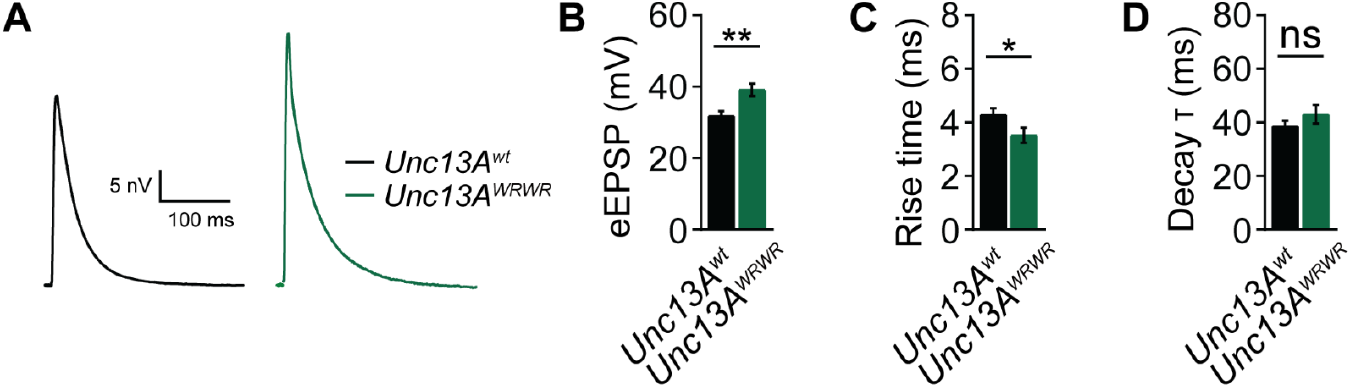
Unc13A^WRWR^ promotes synaptic transmission in *wt* background at larval neuromuscular junction. Related to Figure 2. **(A-D)** Current clamp analysis at 0.4 mM extracellular Ca^2+^ level for *Unc13A*^*WRWR*^ compared to *Unc13A*^*wt*^, including representative evoked excitatory postsynaptic potential (eEPSP) traces (**A**), eEPSP amplitude (**B**), rise time (**C**) and decay constants (**D**). n = 10. *p < 0.05; **p < 0.01; ns, not significant. Error bars: mean ± SEM.

**Supplementary Figure S2.**
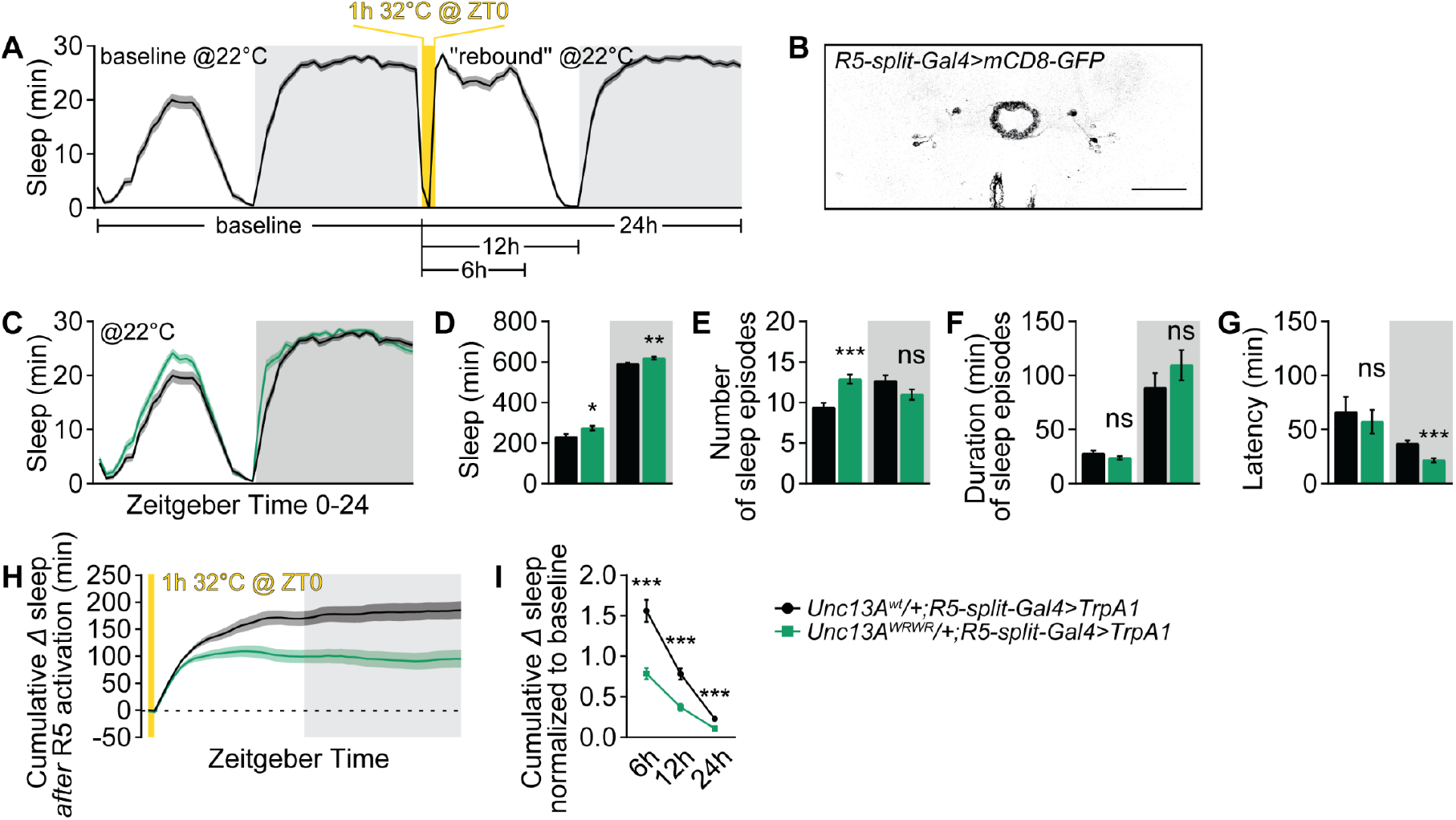
Unc13A^WRWR^/+ suppresses R5 neurons-mediated rebound-like sleep. Related to Figure 2. (**A** and **B**) Averaged sleep curve (**A**) depicting the protocol of inducing rebound-like sleep by briefly and thermogenetically activating ellipsoid body R5 neurons (**B**). (**C**-**G**) Baseline sleep pattern of *Unc13A*^*wt*^/+;*R5-split-Gal4>TrpA1* and *Unc13A*^*WRWR*^*/+;R5-split-Gal4>TrpA1* flies averaged from measurements over 2 days, including sleep profile plotted in 30-min bins (**C**), daytime and nighttime sleep amount (**D**), number and duration of sleep episodes (**E** and **F**), and sleep latency at ZT0 or ZT12 (**G**). n = 94-96. (**H** and **I**) Cumulative gain of sleep during and after the activation of R5 neurons (**H**), and cumulative gain of sleep normalized to baseline sleep within 6, 12 and 24 h since the start of activating R5 neurons (**I**). n = 94-96. *p < 0.05; **p < 0.01; ***p < 0.001; ns, not significant. Error bars: mean ± SEM.

**Supplementary Figure S3.**
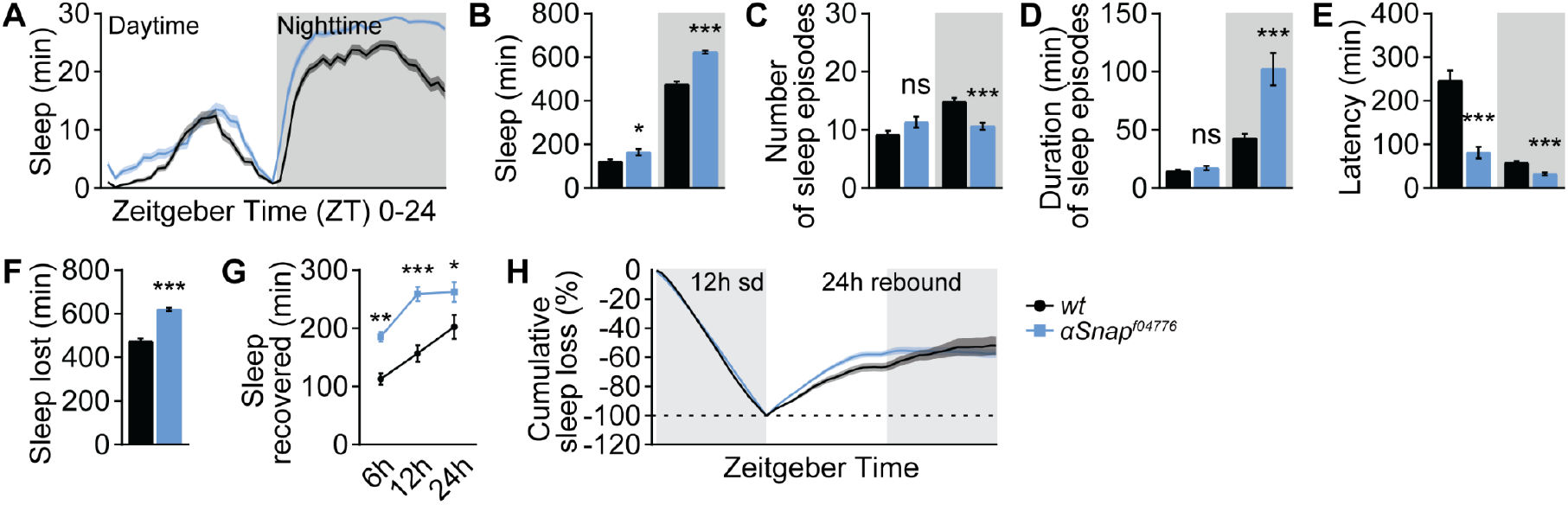
*α*Snap suppresses both baseline and rebound sleep. Related to Figure 3. (**A-E**) Baseline sleep pattern of *wt* and *αSnap*^*f04776*^ flies in *wt* background averaged from measurements over 2 days, including sleep profile plotted in 30-min bins (**A**), daytime and nighttime sleep amount (**B**), number and duration of sleep episodes (**C** and **D**), and sleep latency at ZT0 or ZT12 (**E**). n = 68-70. (**F-H**) Sleep deprivation and sleep rebound analysis including sleep loss due to nighttime sleep deprivation (**F**), absolute recovered sleep within 6, 12 and 24 h after sleep deprivation (**G**), and normalized cumulative sleep loss during 12 h nighttime sleep deprivation and subsequent 24 h sleep rebound (**H**). sd, sleep deprivation. n = 68-70. *p < 0.05; **p < 0.01; ***p < 0.001; ns, not significant. Error bars: mean ± SEM.

